# Defining the parameters for sorting of different RNA cargo into Extracellular vesicles

**DOI:** 10.1101/2024.11.20.624351

**Authors:** Ahmed Abdelgawad, Yiyao Huang, Olesia Gololobova, Yanbao Yu, Kenneth W. Witwer, Vijay Parashar, Mona Batish

**Affiliations:** Department of Biological Sciences, University of Delaware, Newark, DE, USA; Department of Medical and Molecular Sciences, University of Delaware, Newark, DE, USA; Department of Molecular and Comparative Pathobiology, Johns Hopkins University School of Medicine, Baltimore, MD, USA; Department of Laboratory Medicine, Nanfang Hospital, Southern Medical University, Guangzhou, Guangdong, China; Department of Chemistry and Biochemistry, University of Delaware, Newark, DE, USA; Department of Neurology, Johns Hopkins University School of Medicine, Baltimore, MD, USA; Richman Family Precision Medicine Center of Excellence in Alzheimer’s Disease, Johns Hopkins University School of Medicine, Baltimore, MD, USA

**Keywords:** extracellular vesicles, exosomes, circRNAs, mRNAs, lncRNAs, enriched, cis elements

## Abstract

Extracellular vesicles (EVs) are small particles that are released by cells and mediate cell-cell communication by transferring bioactive molecules. RNA cargo of EVs, including coding and non-coding RNAs, can change the behavior of recipient cells, affecting processes like gene expression, proliferation, and apoptosis. Circular RNAs (CircRNA) are a newly appreciated class of regulatory RNAs that are stable, resistant to degradation and have been shown to be enriched in EVs. They play key roles in gene regulation and are also emerging as promising biomarkers for disease diagnosis. While microRNAs (miRNAs) are the most well studied RNA cargo of EVs, very little is known about the mechanisms of enrichment of circRNAs as well as long linear RNAs. Here, we take a comprehensive genome-wide approach to investigate the role of GC%, size, exon count, structuredness and coding potential, in the sorting and enrichment of circular and long linear RNAs into EVs. We found that size and structuredness had a significant role in enriching RNAs into EVs which was consistent across all classes of RNAs. Furthermore, we found that structuredness could explain the relative enrichment of circRNAs over their linear counterparts. These results were validated on existing public databases of circular and linear RNAs in EVs. By identifying and analyzing these factors, we aim to better understand the complex mechanisms behind EV-mediated RNA transfer and its impact on cell communication in both health and disease. This mechanistic understanding of RNA enrichment in EVs is crucial for engineering EVs with selective RNA cargo.

## 1. Introduction

Extracellular vesicles (EVs) are small membrane-bound particles that are secreted by cells into the extracellular environment and play a crucial role in intercellular communication (Gurung et al., 2021; Yáñez-Mó et al., 2015). EVs transfer bioactive molecules, including proteins, and various types of RNAs, between cells and thereby influencing recipient cell behavior (Pitt et al., 2016). Among the diverse cargo of EVs, RNA molecules have garnered significant interest due to their potential in regulating gene expression in recipient cells (Prieto-Vila et al., 2021). The RNA cargo in EVs is biologically active and can significantly alter the behavior of recipient cells (Valadi et al., 2007). For instance, microRNAs (miRNAs) delivered by EVs can suppress the expression of target genes by binding to complementary mRNA sequences, leading to gene silencing (Valadi et al., 2007; Ong et al., 2014; Ding et al., 2015; Viñas et al., 2016). This process can influence cellular functions in recipient cells such as proliferation, differentiation, and apoptosis (He et al., 2020; Ding et al., 2021; Raimondo et al., 2020). The selective enrichment of certain RNA species in EVs suggests that RNA packaging into EVs is a highly regulated process rather than a random event. Most research on RNA sorting into EVs has focused on small non-coding RNAs, primarily miRNAs. RNA-binding proteins like Y-box-binding protein 1 (YBX1), Heterogeneous nuclear ribonucleoprotein A2/B1 (hnRNP A2/B1), and synaptotagmin-binding cytoplasmic RNA-interacting protein (SYNCRIP) have been postulated to assist in sorting miRNAs by binding to small motifs called “zip codes” within the RNA sequence (Liu et al., 2021; Shurtleff et al., 2016; Villarroya-Beltri et al., 2013; Santangelo et al., 2016). A comprehensive review by Leidal and Debnath (2020) further elaborates on these miRNA sorting mechanisms into EVs, highlighting the complexity of this process (Leidal and Debnath, 2020). However, recent work has suggested that the enrichment of miRNAs in EVs might have been overstated, and that the majority of miRNAs reported as EV cargo could simply be part of the EV bio corona (Albanese et al., 2021). On the other hand, the mechanisms underlying the sorting and selection of longer linear RNAs, such as long non-coding RNAs (lncRNAs), and circular RNAs (circRNAs) into EVs remain largely unknown. These RNA species have recently been appreciated to be highly enriched in EVs (Kenneweg et al., 2019; Li et al., 2015), prompting further investigation into their sorting mechanisms.

CircRNAs, in particular, have recently garnered significant attention due to their unique structure and high stability as they are resistant to exonucleases (Zhou et al., 2020). CircRNAs are produced by back-splicing where a downstream 5’ splice site attacks an upstream 3’ splice site joining the intervening sequence into a circular form (Kristensen et al., 2019). CircRNAs play a crucial role in regulating cellular gene expression in various ways. An in-depth review by Kristensen et al. highlights the diverse functions of circRNAs, including their ability to act as sponges for miRNAs by binding them and preventing their interaction with target mRNAs, thus modulating gene expression. Additionally, circRNAs can interact with RNA-binding proteins, influencing cellular processes such as transcription and splicing (Kristensen et al., 2019). Furthermore, the expression of circRNA is often dysregulated in many pathological conditions such as cancer, cardiovascular diseases, and neurological disorders (Verduci et al., 2021; Wang et al., 2023). Due to their stability and disease-specific expression, circRNAs are emerging as promising biomarkers for diagnostics (Allegra et al., 2022). Detecting specific circRNAs in body fluids such as blood or urine could help in the early detection of diseases (Lin et al., 2020; Peter et al., 2021; Zhu et al., 2020). Interestingly, circRNAs were reported to be enriched in EVs compared to their cognate linear isoforms (Lasda and Parker, 2016). This enrichment was first identified by Li et al., who discovered over 1000 circRNAs in human serum exosomes and demonstrated their enrichment in liver cancer cell exosomes (Li et al., 2015). While initially considered a passive process for cells to eliminate these stable RNAs, mounting evidence suggests functional roles for EV-bound circRNAs, implying a selective packaging process. For instance, a recent study by Ngue et al. revealed that circRNAs in breast cancer EVs are associated with chemotherapy resistance, highlighting their potential as both biomarkers and therapeutic targets (Ngue et al., 2023). Furthermore, Chen et al. demonstrated the presence and potential roles of circRNAs in bronchoalveolar lavage fluid (BALF) EVs during lung inflammation, expanding our understanding of circRNA functions in disease contexts (Chen et al., 2024). Despite these advances, the mechanisms underlying the sorting and enrichment of circRNAs into EVs remain elusive.

We hypothesize that the EV enrichment of circRNAs is a cumulative effect of multiple cis elements of the RNA molecule. To test this hypothesis, we performed a genome-wide analysis investigating the role of various factors for circRNA enrichment into EVs, including the presence of zip codes, GC content, size, exon count, and coding potential. Some of these factors have been reported individually in previous studies to impact RNA transport but their cumulative effect is not fully explored. Additionally, we evaluated RNA structuredness as a critical novel factor affecting RNA enrichment into EVs. To our knowledge, this is the first study to understand the role of RNA structure on its transport. Furthermore, we also analyzed the role of these factors on the transport of long linear RNAs including mRNAs and lncRNAs. Our study aims to develop a comprehensive model to understand the process of RNA packaging into EVs and provide tools to predict the potential of a given RNA molecule to be enriched in EVs. By identifying and characterizing these factors, we hope to shed light on the intricate processes underlying EV-mediated RNA transfer and their implications for cell-to-cell communication in health and disease. This research has the potential to advance our understanding of EV biology and open new avenues for diagnostic and therapeutic applications leveraging EV-RNA interactions.

## 2. Results

### 2.1. CircRNAs are enriched in EVs

The most well studied RNA cargo of extracellular vesicles (EVs) are the miRNAs which are small regulatory noncoding RNAs (Cheng et al., 2014; Vaka et al., 2023; Honorato-Mauer et al., 2023; Park et al., 2020; Liu et al., 2019). However, recent work has shown that other RNA species including but not limited to mRNAs, lncRNAs and circRNAs are most present in EVs and have been shown to play functional roles (Li et al., 2015; Kenneweg et al., 2019). Despite a lack of clear understanding of the process of RNA cargo selection, several studies have reported that miRNAs get sorted via binding to specific RBPs (Villarroya-Beltri et al., 2013; Chen et al., 2009; Li et al., 2012; Santangelo et al., 2016; Hobor et al., 2018). These RBPs recognize a short sequence in the miRNA (often referred to as “zip-code”) and sort those miRNAs in EVs. Similar “Zipcodes” have not been well established for other EV RNA cargo. We sought to characterize the features of other linear and circular RNAs that are known to be enriched in EVs and test if they harbor similar “Zipcodes” to the ones identified for miRNA enrichment. To get a holistic view, we prepared RNA sequencing libraries of total RNAs from DLD-1 cells and their isolated EVs. EVs from DLD-1 cells were isolated and characterized according to MISEV guidelines (Théry et al., 2018) (Figure 1A-C, Extended Data Figure 1A). Although circRNAs are much more stable compared to linear RNAs—due to their lack of free ends, they represent a small proportion of cellular RNAs (Zhang et al., 2019). To enrich circRNAs, RNA from both cell and EVs fractions were treated with RNase R which digests linear RNAs and, thus, enriches circRNAs. Samples had a high percentage of reads aligning to the human genome (Extended Data Figure 1B). CircRNAs were then identified from sequencing data and analyzed for enrichment in EVs. First, we did observe a significant increase in the ratio of circRNAs to linear RNAs after RNase R treatment in both cellular and EVs fractions (Extended Data Figure 1C). Further, and consistent with previous reports of circRNAs enrichment in EVs, we found that in control samples (no RNase R treatment), the ratio of circular to linear RNAs was significantly higher in EVs compared to cells (0.288 vs 0.178; p < 0.001, Mann-Whitney’s test) (Figure 1D). This suggests that circRNAs are more enriched in EVs compared to the cell which is in line with previous reports (Li et al., 2015; Lasda and Parker, 2016). Furthermore, we compared the raw counts and normalized counts of different classes of RNAs in EVs. Since, using the number of back-splice junctions severely underestimates circRNAs counts, we remapped the reads of RNase R-treated samples to circRNAs detected using back-splice junctions. Comparing the number of transcripts of both circRNAs and linear RNAs (mRNA, lncRNAs and other RNAs) that are detected in EVs, we observed higher average count of circRNAs transcripts compared to linear classes of RNAs (Figure 1E). There were more circRNAs detected in at least one or two of the three replicates compared to linear RNAs. For the RNAs detected in all of the three replicates, mRNAs had slightly higher count followed by circRNAs, others and, finally, lncRNAs. Moreover, RNA counts from all classes were normalized using DESeq2 and their differential enrichment in EVs were analyzed. CircRNAs showed the highest enrichment in EVs followed by mRNAs, lncRNAs, and other RNAs (Figure 1F). These results are again consistent with previous reports and highlight the preferential enrichment of circRNAs in EVs.

**Figure 1.**
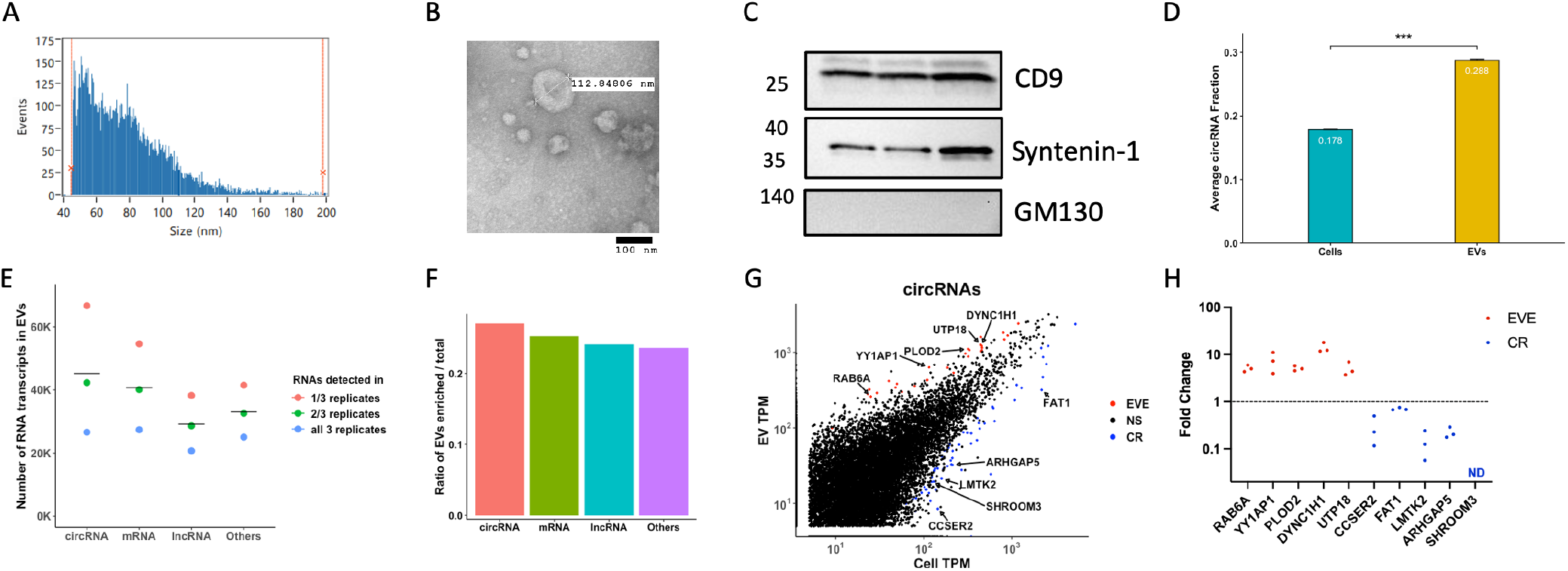
CircRNAs are enriched in EVs: **(A)** Size profiles of isolated EVs by nano-flow cytometry from one representative EV sample. **(B)** representative image of EVs visualized by negative staining transmission electron microscopy (scale bar= 100 nm). **(C)** Western blot of three EVs samples. **(D)** Average circular/linear RNA fraction in cell and EVs. **(E)** Total number of transcripts detected in either 1, 2, or all three EV samples from RNA-Seq for all four classes of RNAs analyzed. **(F)** Ratio of significantly enriched transcripts in EVs compared to cellular fraction of the four classes of RNAs analyzed. **(G)** circRNAs abundance from RNA-Seq in EVs and cells. Arrows point to names of circRNAs that were validated by RT-qPCR. **(H)** EVs – cell fold Change of circRNAs from three independent samples. ND = not detected, EVE = EVs enriched, CR = cell retained. P-values were calculated using Mann-Whitney’s test, *p < 0.05, **p < 0.01, ***p < 0.001.

Overall, we observed a high correlation between the abundance of circRNAs in cells and in EVs (r = 0.84) and we validated selected enriched and depleted circRNAs using RT-qPCR (Figure 1G-H). These results validated that circRNAs are one of the major RNA cargoes in EVs.

### 2.2. Role of cis elements in enriching circRNAs in EVs

We next asked if there is any selection amongst circRNAs regarding which ones will be enriched into EVs and which will be retained in the cell. We wondered if the EVs-enriched circRNAs harbor the same “Zipcode” elements that are reported to facilitate the enrichment of miRNAs in EVs. Towards this end, we compiled a list of five different RBPs that were reported to be involved in RNA transport into EVs and retrieved their motifs from ATtRACT database (Giudice et al., 2016). For all circRNAs in our dataset, we compared the motif occurrence of the five RBPs in each circRNA and compared it to the 3’ UTR of its parent gene. This analysis showed that 3’UTRs contained higher counts of RBPs motifs compared to circRNAs since the average difference in motif counts of circRNAs and 3’UTRs of their counterpart linear transcripts were negative (Figure 2A). The only exception was hnRNPA2B1 which showed a positive, albeit small, value of 0.002, suggesting that hnRNPA2B1 motif counts were almost similar between circRNAs and the 3’ UTRs of their respective parent genes.

**Figure 2.**
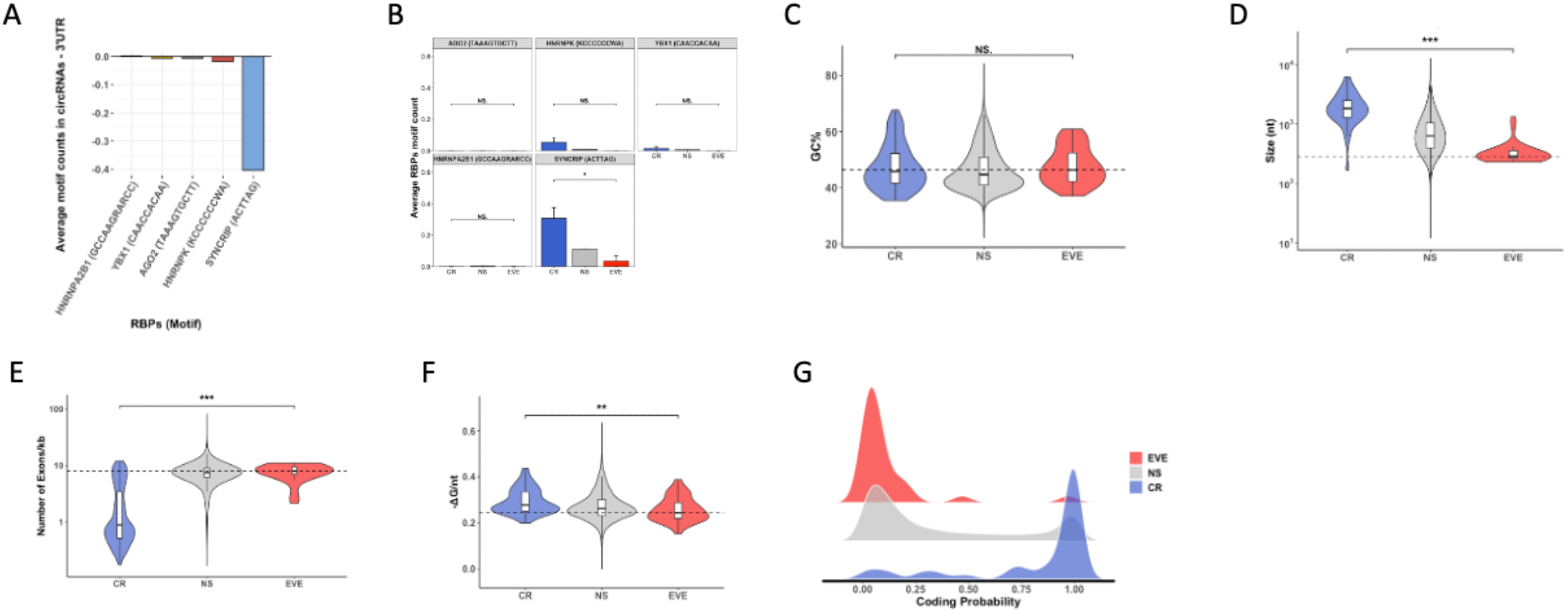
Evaluation of role of cis elements in EVs enrichment of circRNAs: **(A)** Average motif count difference between circRNAs and the 3’ UTR of their linear counterparts for five RBPs. Motifs of RBPs are shown in parenthesis. **(B)** Average motif count of five RBPs of circRNAs that are EVs enriched (EVE), cell retained (CR), or those with no significant difference (NS). Motifs of RBPs are shown in brackets. **(C-F)** Violin plot of GC%, size, number of exons per kilobase of transcript, and structuredness, respectively, of CR, NS, and EVE circRNAs. **(G)** Density plot of coding probability of CR, NS, and EVE circRNAs. EVE = EVs enriched, NS = not significant. CR = cell retained. P-values were calculated using Mann-Whitney’s test, *p < 0.05, **p < 0.01, ***p < 0.001.

Next, within our pool of circRNAs, we categorized them based on their enrichment in EVs as either EVs-enriched (EVE), cell retained (CR) or those whose enrichment is not significant (NS), and we then compared the motif occurrence of all five RBPs in these three categories of circRNAs. All five RBPs either had similar or higher average motif count in CR category compared to either EVE category or those in NS category (Figure 2B). In the case of SYNCRIP, CR circRNAs had significantly higher counts of RBPs motifs compared to EVE circRNAs (mean 0.307 vs 0.034, p = 0.011, Mann-Whitney’s test). Since, size might impact the motif counts in a sequence, we also calculated the fraction of circRNAs containing RBPs motifs. Consistently, CR circRNAs had higher percentage of those containing RBPs motifs (Extended Data Figure 2A). Overall, these results indicated that circRNA enrichment over linear RNAs in EVs could not be solely explained by the presence of “Zipcodes”. A recent study has investigated the role of other cis elements including size, GC%, and number of exons in sorting linear RNAs into EVs (O’Grady et al., 2022). This prompted us to investigate the role of these cis elements in circRNAs enrichment in EVs.

For GC%, we did not observe a significant difference in between EVE and CR circRNAs (median 46.375 vs 45.859, p = 0.572, Mann-Whitney’s test) (Figure 2C). For size, we found EVE circRNAs to be significantly smaller than CR circRNAs (median 285 vs 1832, p = 3.214×10^−13^, Mann-Whitney’s test) (Figure 2D). Interestingly, despite having smaller size, EVE circRNAs had significantly higher number of exons per kilobase of transcript (kb) compared to CR circRNAs (median 7.968 vs 0.889, p = 8.849×10^−10^, Mann-Whitney’s test) (Figure 2E). However, exons in EVE circRNAs were on average smaller than those in CR circRNAs (135 nt vs 569 nt). Together, these data suggest that circRNAs that are enriched in EVs are smaller in size and has more exons relative to their size.

Most of the studies explore the features of primary RNA sequence but often overlook the role of secondary structure of the RNA. The secondary structure of RNA was recently shown to be involved in RNA decay where highly structured RNAs are more likely to be degraded as compared to low structured RNAs (Fischer et al., 2020). We hypothesized that since low structured RNAs do not get targeted for degradation, they will be more likely to get packaged into EVs. To test this hypothesis. we calculated the degree of structuredness of all circRNAs in our dataset using RNAfold’s size-normalized minimum free energy (-ΔG/nt) (Lorenz et al., 2011). This algorithm showed high correlation with experimental methods of determining RNA secondary structure such as DMS-Seq (Fischer et al., 2020). Indeed, we found that EVE circRNAs showed a significantly lower structuredness compared to CR circRNAs (median 0.244 vs 0.277, p = 0.003, Mann-Whitney’s test) (Figure 2F).

Finally, since some of the circRNAs have been reported to be translated into proteins, we also sought to see if the coding potential of circRNAs in the cell would affect their EVs enrichment (Legnini et al., 2017; Jiang et al., 2021; van Heesch et al., 2019). To address this question, we utilized CPC2, an algorithm that calculates the coding potential of RNAs based on sequence features (Kang et al., 2017). We found that EVE circRNAs had noticeably lower coding potential than CR circRNAs (median 0.052 vs 0.995, p = 4.12×10^−12^, Mann-Whitney’s test) (Figure 2G). Together, these results indicate that EVE circRNAs have lower structuredness and are less likely to be translated into proteins.

### 2.3. CircRNA validation using public databases

Since, we have used RNA samples from cells and EVs of a single cell line, we wanted to verify if these findings will also be applicable to a larger set of RNAs from diverse sources. Therefore, we evaluated two publicly available databases including circBase, which catalogs circRNAs from diverse sources and ExoRBase, a database containing circRNAs reported in human blood EVs (Glažar et al., 2014; Lai et al., 2022). We utilized detection frequency as a proxy for EVs enrichment and arbitrarily chose a cutoff threshold of 0.5 to compare circRNAs with high and low detection frequency in ExoRBase. We also compared circRNAs in ExoRBase to those in CircBase as a control after removing CircBase circRNAs that are included in exoRBase.

First, we analyzed circRNAs in both exoRBase and circBase and calculated the difference in RBPs motif counts between circRNAs and the 3’ UTR of their respective genes. Results were mostly similar to that from our dataset where most RBPs had lower average counts in circRNAs compared to their linear counterparts in both datasets (Figure 3A and Extended Data Figure 3A). Next, we analyzed circRNAs for GC%. Consistent with our data, circRNAs in exoRBase with high detection frequency showed no significant difference than those with low detection frequency (median 43.269 vs 43.589, p = 0.31, Mann-Whitney’s test) (Figure 3B). However, circRNAs in ExoRBase showed significantly lower GC% than those in CircBase (median 43.584 vs 47.452, p < 2.2×10^−16^, Mann-Whitney’s test) (Extended Data Figure 3B).

**Figure 3.**
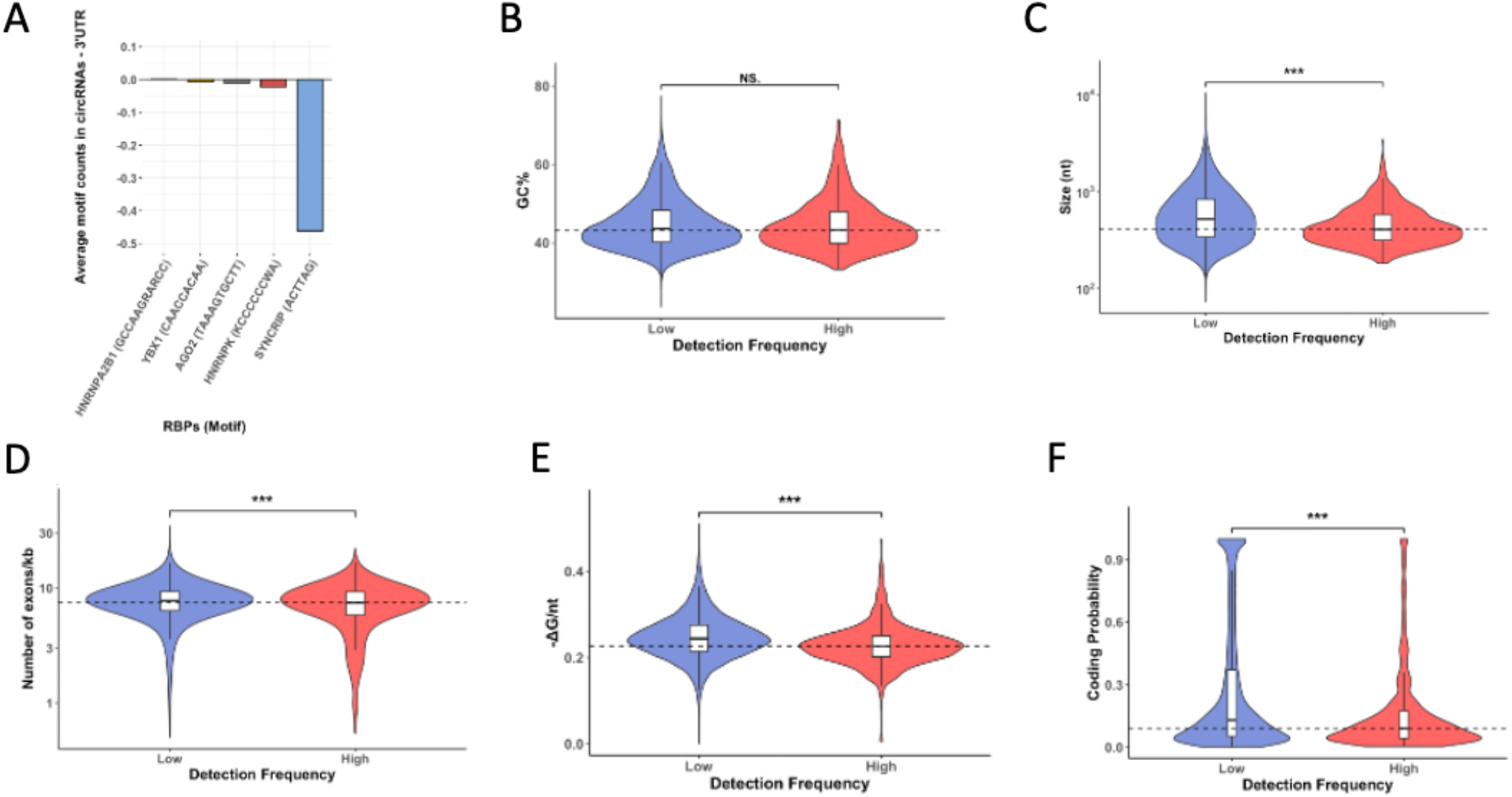
Validation of circRNAs cis elements from databases: **(A)** Average motif count difference between circRNAs in exoRBase and the 3’ UTR of their linear counterparts for five RBPs. Motifs of RBPs are shown in parenthesis. **(B-F)** Violin plot of GC%, size, number of exons per kilobase of transcript, structuredness, and coding probability, respectively, of exoRBase circRNAs with low or high detection frequency. P-values were calculated using Mann-Whitney’s test, *p < 0.05, **p < 0.01, ***p < 0.001.

Consistently with the results from our dataset, the analysis of size in exoRBase showed that circRNAs with high detection frequency were significantly smaller than those with low detection frequency (median 406 vs 520, p < 2.2×10^−16^, Mann-Whitney’s test) (Figure 3C). Similarly, exoRBase circRNAs were significantly smaller than those in CircBase (median 518 vs 710, p < 2.2×10^−16^, Mann-Whitney’s test) (Extended data Figure 3C). The number of exons per kilobase of transcript of circRNAs in exoRBase was significantly higher than circRNAs in circBase (median 7.752 vs 7.247, p < 2.2×10^−16^, Mann-Whitney’s test) (Extended data Figure 3D). However, within exoRBase, the number of exons per kilobase of transcript was significantly lower in circRNAs with high detection frequency than those with low detection frequency, albeit by a smaller margin (median 7.463 vs 7.752, p < 0.0007, Mann-Whitney’s test) (Figure 3D). Next, we compared the structuredness (-ΔG/nt) of circRNAs. CircRNAs from exoRBase with high detection frequency had significantly lower structuredness compared to those with low detection frequency (median 0.226 vs 0.244, p < 2.2×10^−16^, Mann-Whitney’s test) (Figure 3E). Similarly, ExoRBase circRNAs had significantly lower structuredness compared to those in circBase (median 0.244 vs 0.276, p < 2.2×10^−16^, Mann-Whitney’s test) (Extended data Figure 3E). Further, we compared the coding probability of circRNAs. Again, both comparisons showed consistent results with our data. For ExoRBase, circRNAs with high detection frequency showed significantly lower coding probability than those with low detection frequency (0.09 vs 0.131, p < 1.095×10^−14^, Mann-Whitney’s test) (Figure 3F). Similarly, CircRNAs in ExoRBase had significantly lower coding probability than those in circBase (median 0.13 vs 0.216, p < 2.2×10^−16^, Mann-Whitney’s test) (Extended data Figure 3F). Overall, these results highlight that most cis elements compared using published databases showed consistent results with our data which increases confidence in our findings

### 2.4. Role of cis elements in enriching linear RNAs in EVs

Next, we investigated if the above studied factors are also involved in the EVs enrichment of linear RNAs. To this end, we first separated all linear RNAs from Gencode v32 into three categories: protein-coding genes, labeled as mRNAs, long non-coding RNAs labeled as lncRNAs and all other classes of linear RNAs labeled as others. We then quantified linear RNAs in our RNA-Seq data and categorized them into EVE, NS, or CR based on their enrichment in EVs. We used qPCR to validate select mRNA and lncRNA candidates from both EVE and CR categories and obtained consistent results to our RNA-Seq datasets (Figure 4A-D).

**Figure 4.**
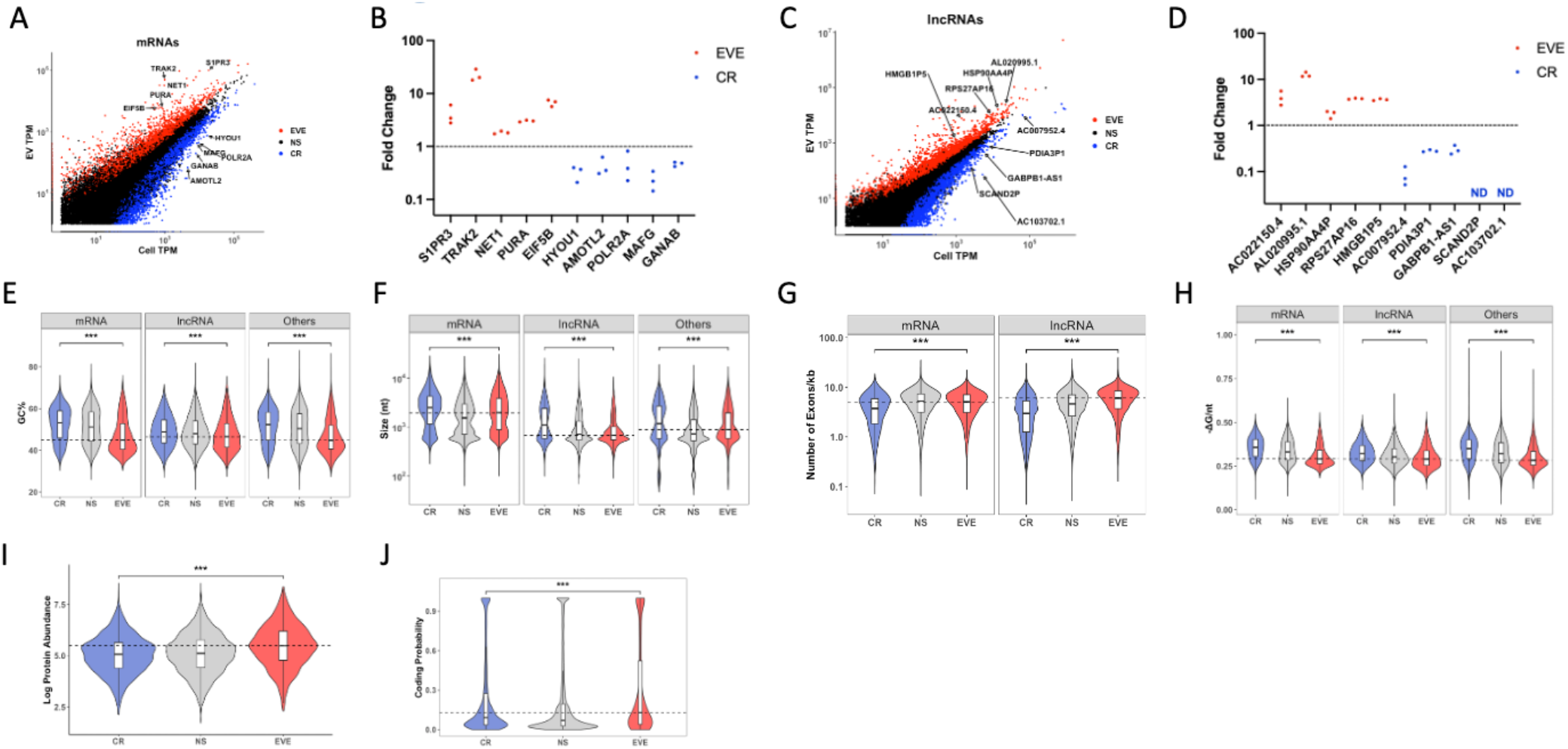
Evaluation of role of cis elements in EVs enrichment of linear RNAs: **(A)** mRNAs abundance from RNA-Seq in EVs and cells. Arrows point to names of mRNAs that were validated by RT-qPCR. **(B)** EVs – cell fold Change of mRNAs from three independent samples. ND = not detected. **(C-D)** Similar to A, B but for lncRNAs. **(E)** Violin plot of GC% in mRNAs, lncRNAs, and others separated based on their enrichment into EVE, NS and CR RNAs. **(F)** Similar to R but for size. **(G)** Violin plot of number of exons per kilobase of transcript in mRNAs and lncRNAs separated based on their enrichment into EVE, NS, and CR RNAs. **(H)** Similar to E but for structuredness. **(I)** Log protein cellular abundance of EVE, NS, and CR mRNAs. **(J)** Violin plot of coding probability of EVE, NS and CR lncRNAs. EVE = EVs enriched, NS = not significant. CR = cell retained. P-values were calculated using Mann-Whitney’s test, *p < 0.05, **p < 0.01, ***p < 0.001.

To determine the role of RNA features identified for circRNAs in linear RNAs, we first evaluated the GC% of three classes of liner RNAs. EVE RNAs from all three categories had lower GC% compared to CR RNAs (median for mRNAs: 44.865 vs 53.09, p < 2.2×10^−16^; median for lncRNAs: 46.52 vs 48.75, p < 2.2×10^−16^; median for Others: 44.75 vs 52.35, p < 2.2×10^−16^, Mann-Whitney’s test) (Figure 4E). We further dissected mRNAs into their constituting parts: 5’ UTR, CDS and 3’UTR. Both CDS and 3’UTR. We found the GC% of both CDS and 3’ UTR were significantly lower in EVE mRNAs compared to CR mRNAs (median for CDS: 45.87 vs 55.61, p < 2.2×10^−16^; median for 3’UTR: 35.9 vs 48.125, p < 2.2×10^−16^, Mann-Whitney’s test), whereas 5’UTR GC% was higher in EVE mRNAs compared to CR mRNAs (median 63.5 vs 61.93, p = 3.5×10^−7^, Mann-Whitney’s test) (Extended data Figure 4A).

For size, all three RNA classes showed significantly lower size for EVE RNAs compared to CR RNAs (median for mRNAs: 1964 vs 2487, p < 2.2×10^−16^; median for lncRNAs: 678 vs 1096, p < 2.2×10^−16^; median for Others: 884 vs 1167, p < 2.2×10^−16^, Mann-Whitney’s test) (Figure 4F). Furthermore, all parts of mRNAs showed a significantly smaller size for EVE mRNAs compared to CR mRNAs with 3’UTR showing the most significant difference (median for 5’ UTR: 164 vs 187, p = 2.266×10^−14^; median for CDS: 1008 vs 1041, p = 0.01; median for 3’ UTR: 652 vs 1014, p < 2.2×10^−16^, Mann-Whitney’s test). This was consistent with size comparison in circRNAs and suggests that physical size limitation of EVs could be a factor in selecting cargo for EVs (Extended data Figure 4B).

Interestingly, for mRNAs, we found EVE mRNAs to have significantly more exons per kilobase of transcript compared to CR mRNAs (median 5.066 vs 3.738, p < 2.2×10^−16^, Mann-Whitney’s test). However, these exons in EVE mRNAs were on average smaller than those in CR mRNAs (246.413 nt vs 336.112 nt) (Figure 4G). Similarly, lncRNAs showed a similar difference between EVE and CR lncRNAs (mean 6.038 vs 2.967, p < 2.2×10^−16^, Mann-Whitney’s test) and EVE lncRNAs exons were also smaller on average compared to those in CR lncRNAs (207.923 vs 481.1427) (Figure 4G).

Next, we investigated structuredness of linear RNAs. Interestingly, all classes of RNAs that are enriched in EVs showed significantly lower structuredness than those retained in the cell (median for mRNAs: 0.291 vs 0.356, p < 2.2×10^−16^; median for lncRNAs: 0.289 vs 0.322, p < 2.2×10^−16^; median for others, 0.283 vs 0.35, p < 2.2×10^−16^, Mann-Whitney’s test) (Figure 4H). Next, for mRNAs, most of the difference was attributed to 3’ UTR which showed the biggest difference in structuredness (median for 3’ UTR of EVE mRNAs 0.228 vs 0.312, p < 2.2×10^−16^, Mann-Whitney’s test). CDS of EVE mRNAs was also significantly less structured compared to CDS of CR mRNAs (median 0.281 vs 0.348, p < 2.2×10^−16^, Mann-Whitney’s test). However, 5’UTR structuredness of EVE mRNAs was not significantly different from that of CR mRNAs (0.352 vs 0.349, p = 0.107, Mann-Whitney’s test) (Extended data Figure 4C). This data is once again consistent with circRNAs and highlights the universality of structuredness as a factor in selecting RNA cargo in EVs.

Finally, we looked at the coding probability of lncRNAs. In our dataset, EVE lncRNAs showed higher coding probability compared to CR lncRNAs (median 0.1128 vs 0.092, p < 2.2×10^−16^, Mann-Whitney’s test) (Figure 4I). For mRNAs, we sought to correlate RNA abundance in EVs with proteins abundance in the cell, we performed mass spectrometry analysis of DLD-1 whole-cell lysate. We then correlated transcript per million (TPM) values for mRNAs with intensity-based absolute quantification (iBAQ) for proteins (Schwanhäusser et al., 2011). Surprisingly, Spearman’s correlation of protein abundance with RNA abundance in EVs was higher than the correlation with RNA abundance in cells (0.487 vs 0.457, p-value = 0.002). We also saw a significantly higher protein abundance of EVE mRNAs compared to CR mRNAs (median 5.413 vs 4.992, p < 2.2×10^−16^, Mann-Whitney’s test) which is consistent with lncRNAs (Figure 4J). These data suggest that coding potential of RNAs might have opposing effect for circular and linear RNAs enrichment in EVs.

### 2.5. Linear RNA validation using public databases

Similar to our extended analysis for circRNAs, we sought to confirm our findings for linear RNAs using publicly available datasets. We analyzed mRNAs and lncRNAs from exoRBase and utilized detection frequency as a proxy for EVs enrichment with a cutoff threshold of 0.75 and 0.5 for mRNAs and lncRNAs, respectively.

For GC%, mRNAs with high detection frequency in ExoRBase had lower GC% compared to those with low detection frequency (median 50.7 vs 53.3, p < 2.2×10^−16^, Mann-Whitney’s test) (Figure 5A). However, lncRNAs with high detection frequency had significantly higher GC% (median 49.505 vs 47.57, p = 7.95×10^−11^, Mann-Whitney’s test) which is contrary to our dataset where we found EVE lncRNAs to have lower GC% (Extended data Figure 5A).

**Figure 5.**
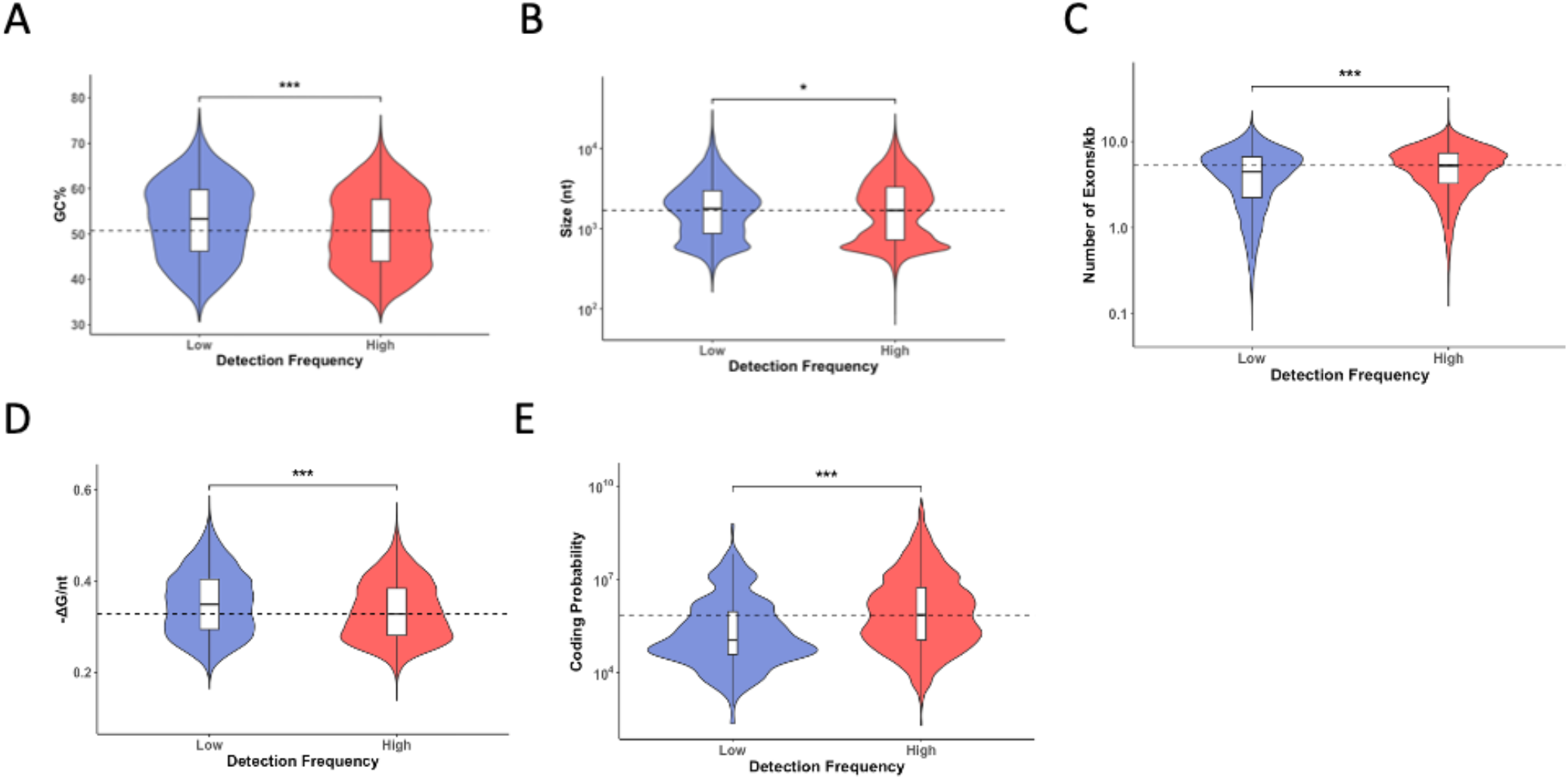
Validation of mRNAs cis elements from databases: **(A-E)** Violin plot of GC%, size, number of exons per kilobase of transcript, structuredness, and coding probability, respectively, of exoRBase mRNAs with low or high detection frequency. P-values were calculated using Mann-Whitney’s test, *p < 0.05, **p < 0.01, ***p < 0.001.

For size, mRNAs with high detection frequency had significantly smaller size compared to those with low detection frequency (median 1697 vs 1773, p = 0.018, Mann-Whitney’s test). LncRNAs showed inconsistent results with our data with high detection-frequency lncRNAs having significantly bigger size than those with low detection frequency (median 802 vs 635, p < 2.2×10^−16^, Mann-Whitney’s test) (Extended data Figure 5B).

High detection-frequency mRNAs had consistently more exons per kilobase of transcript compared to those with low detection frequency (median 5.261 vs 4.464, p < 2.2×10^−16^, Mann-Whitney’s test) (Figure 5C). However, number of exons per kilobase of transcript was not significantly different between lncRNAs with high detection frequency and those with lower detection frequency (median 3.484 vs 3.521, p < 0.162 Mann-Whitney’s test) (Extended data Figure 5C).

High detection-frequency mRNAs were also significantly less structured compared to those with low detection frequency (median 0.328 vs 0.349, p < 2.2×10^−16^, Mann-Whitney’s test) (Figure 5D). LncRNAs in ExoRBase, however, did not show similar results to our datasets where lncRNAs with high detection frequency showed higher structuredness than those with low detection frequency (median 0.317 vs 0.298, p < 2.489×10^−15^, Mann-Whitney’s test) (Extended data Figure 5D).

Finally, estimating the protein abundance of ExoRBase mRNAs with high detection frequency from our dataset, we found that they had significantly higher protein abundance compared to those with lower detection frequency (median 7.082×10^5^ vs 1.111×10^5^, p < 2.489×10^−15^, Mann-Whitney’s test) (Figure 5E). The coding probability of lncRNAs with high detection frequency was significantly higher than that of low detection-frequency lncRNAs (median 0.063 vs 0.047, p < 4.179×10^−10^, Mann-Whitney’s test) (Extended data Figure 5E). Overall, these data suggest good consistency with our data for mRNAs in database whereas lncRNAs in database showed less agreement with results from our dataset.

### 2.6. Circularization enhances RNAs enrichment in EVs

Since, a given gene can be spliced differently to produce both linear RNAs as well as circRNAs and thus linear RNAs harbor most of the same cis elements that are found in circRNAs. We wondered if the linear counterparts of EV-enriched circRNAs will have an equal probability to be enriched into EVs and vice versa. We compared the abundance of linear cognates of circRNAs and found a Spearman correlation of 0.007 indicating no correlation between the enrichment of linear and circular transcripts from the same gene into EVs. Furthermore, we analyzed the differential EV-enrichment of linear isoforms of all circRNAs in our dataset. The distributions of the fold change of linear counterparts were almost identical regardless of the level of enrichment of the circRNAs (Figure 6A). These data further confirm that there is a distinct and independent mechanism for the enrichment of circular and linear RNAs into EVs despite sharing the same sequence.

**Figure 6.**
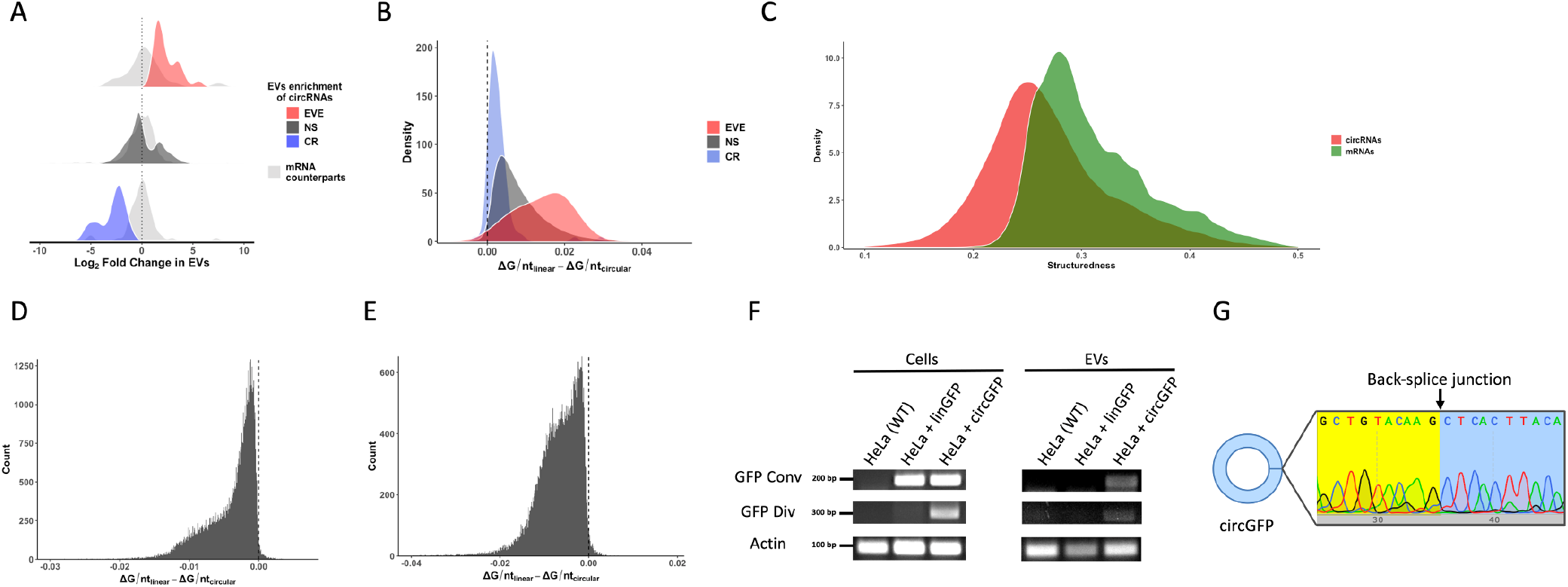
Role of structuredness in EVs enrichment of circRNAs: **(A)** Density plot of log2 fold change in EVs of circRNAs that are EVs enriched (EVE), cell retained (CR), or those with no significant difference (NS) as well as their respective mRNA counterparts. **(B)** Change in structuredness after linearization of EVE, NS, or CR circRNAs sequences. **(C)** Density plot of the structuredness of circRNAs and their mRNA counterparts. **(D-E)** Change in structuredness of after circularization of mRNAs and lncRNAs sequences, respectively. **(F)** Gel image of WT HeLa cells, or cells expressing linear GFP or circular GFP (circGFP) in cells and EVs using both convergent and divergent primers. **(G)** Sequence of circGFP covering the back-splice junction. EVE = EVs enriched, NS = not significant. CR = cell retained.

We then asked if circular conformation could explain circRNAs preferential enrichment in EVs. To address this question, we performed in-silico linearization of circRNAs sequences and calculated the difference in their structuredness scores and compared that to the original scores. It turns out most circRNAs from all three datasets gained structuredness (-ΔG/nt) after linearization (Figure 6B). In our dataset, 98.3% of circRNAs had lower structuredness compared to their linearized sequences. Furthermore, we found larger gains in structuredness after in silico linearization for EVs-enriched circRNAs than those with no enrichment or cell-retained circRNAs (Figure 6B). Linearization of circRNAs from both exoRBase and circBase, showed similar results (Extended data Figure 6A-B). Furthermore, when we compared the structuredness of circRNAs to that of their linear counterparts, we observed that circRNAs, on average, had significantly lower size-normalized structuredness than their linear counterparts (Figure 6C).

We also investigated whether linear RNAs would become less structured after circularization. Circularizing mRNA sequences from both our dataset and from exoRBase, led to lower structuredness (Figure 6D, Extended data Figure 6C). Similarly, lncRNAs showed lower structuredness when their sequences were circularized in both our own dataset as well as in exoRBase (Figure 6E, Extended data Figure 6D). These data suggest that circular conformation leads to lower structuredness of circRNAs which could explain their EVs enrichment over their linear counterparts. To experimentally verify these in silico results, we designed a linear and circular version of GFP, and we cloned either linear GFP or circular form of GFP (circGFP) in HeLa cells and isolated RNAs from both cellular and EV fractions of cells expressing these GFP variants. We found that circGFP could be detected in EVs but not linear GFP (Figure 6F). To confirm the circularity of the circGFP, the PCR products from both cellular and EV fractions were extracted and sequenced to confirm the presence of the back-splice junction (Figure 6G). These results validated our findings that circularization of an RNA could increase its enrichment in EVs likely by lowering its structuredness.

## 3. Discussion

In this study, we evaluated the role of several cis factors in the enrichment of different RNA species into EVs. Our findings suggest that RNA packaging into EVs is dependent on a combination of RNA cis elements including GC%, size, the number of exons, structuredness and the coding potential. While numerous studies have investigated the role of GC%, size and exons count in RNAs enrichment in EVs, this is the first study to explore the role of structuredness and coding potential of RNA in its extracellular transport as well look at combinatorial effect of multiple cis factors. We used a genome wide approach to characterize all the long RNA cargo of EVs. Our data validated the fact that circRNAs are more abundant in EVs compared to other long linear classes of RNA, such as mRNAs and lncRNAs which was also reported by other studies (Li et al., 2015; Lasda and Parker, 2016). The process of EVs cargo selection is likely a highly regulated one where the cellular function and homeostasis of a given RNA could affect its enrichment in EVs. For example, CDR1as, a circRNA well known for its sponging function of miR-7, was not enriched in EVs when miR-7 mimics were introduced in the cell while its cellular levels increased (Li et al., 2015). On the other hand, RNA cargo selection for EVs could potentially be a yet another tool in the arsenal of cell to regulate its gene expression (O’Grady et al., 2022).

The most common mechanism explored for RNA transport thus far relies on the ability of RNA binding proteins to recognize specific sequence motifs in the RNA sequences. This RBP-dependent RNA transport is studied both in the context of the transport of different RNA species within the cell as well for its extracellular Transport (Batish et al., 2012; Fabbiano et al., 2020). For instance, it was believed that neuronal RNAs are transported to distal dendrites via packaging into RNA granules using RBPs by virtue of having the RBP recognition motifs termed as “zip codes”. However, it was shown that addition of zip codes from a dendritically localized RNA to an exogenous RNA does not result in its packaging into RNA granules. In fact, single molecule imaging revealed that RNA granules do not have aggregates of RNAs of same or different species implying that these zip codes do not direct the packaging of RNAs into granules (Batish et al., 2012). Similarly, several zip codes have been identified for EV bound RNAs particularly for miRNAs. We evaluated the presence of these zip codes in EV-enriched circRNAs and long linear RNAs and could not find any significant correlation between the presence of specific zip codes in RNAs and their EVs enrichment. These results signify that the presence of RBP recognition motifs on RNAs cannot solely dictate the packaging of a given RNA species into EVs, and that certain constraints on RNA transport apply in conjunction.

Recent work has identified the correlation of other RNA cis factors with EVs enrichment. Of note, size and GC content are one of most evaluated features. CircRNAs in EVs were also reported to have smaller size and lower GC% compared to cellular circRNAs (Zhang et al., 2019). Consistently, we observed that circRNAs enriched in EVs in our dataset as well as in public databases have a smaller size. Further, GC% of circRNAs with high detection frequency in exoRBase was significantly different. In our dataset, although the GC% difference was not significant, it was slightly lower than that of cell-retained circRNAs. Given the small size of EVs, it is perhaps reasonable to assume that smaller RNAs would have a higher tendency to be packaged as compared to longer RNAs. Furthermore, O’Grady et al., have recently examined the correlation of cis elements such as GC%, size and exon counts with EVs enrichment of linear RNAs and found that EV enriched linear RNAs tend to have smaller size, contain a larger number of exons and have relatively higher GC% (O’Grady et al., 2022). We found similar results for the size and exon count for mRNAs and lncRNAs, but our data suggest that linear RNAs that are EVs enriched have lower GC% compared to those retained in the cell. Some studies, however, have found no difference in GC content of long RNAs between EVs and human plasma while others reported lower GC% of circRNAs in EVs fractions compared to cellular fractions in HepG2 cells (Rodosthenous et al., 2020; Zhang et al., 2019). This inconsistency could be attributed to different cell types used in those studies and, therefore, might suggest an inconsequential role of GC content in enriching RNAs in EVs. The coding potential is a relatively less explored feature of RNA that we showed to have a role in EV packaging of RNA. Interestingly, we found that coding potential has opposite effects for circular and linear RNAs. Analysis of RNAs from public datasets confirmed the effect we observed in our own data. EVs were enriched with circRNAs of lower coding potential and lncRNAs of higher coding potential than their respective cellular fractions. While some circRNAs were recently reported to encode peptides, they still represent a small fraction of identified circRNAs. For instance, about 1,200 circRNAs are predicted to have the ability to be translated into peptides (Li et al., 2021). This number diminishes in comparison to over 90,000 circRNAs reported in circBase and over 289,000 reported in more recent databases such as CircNet 2.0 (Glažar et al., 2014; Chen et al., 2022). Conversely, regulatory functions of circRNAs including miRNAs sponging and protein interactions remain the main functions reported for circRNAs. For instance, the number of predicted circRNA-miRNA interactions are over nine million (Chen et al., 2022). On the other hand, Almeida et al. have reported numerous lncRNAs in EVs that were predicted to encode peptides and have experimentally confirmed several of those peptides (Almeida et al., 2022). Perhaps, cells prioritize circRNAs with non-coding functions and lncRNAs with translation capabilities to be transported into EVs. Furthermore, we quantified the correlation of protein abundance with mRNAs abundance cells and obtained similar results to what is reported in literature (Jarnuczak et al., 2021). It is intriguing to see a small, albeit significant, increase in correlation of protein abundance with mRNAs abundance in EVs compared to cells. Our data also indicate the enrichment of mRNAs that had higher cellular protein abundance. It is possible that cells might opt to restrict the transport of mRNAs with low protein abundance as their function could be needed in the cell.

One of the key findings of this study is the role of structuredness on RNA transport. Our data suggest that structuredness is a novel factor that could explain the preferential enrichment of circRNAs in EVs. RNA secondary structure was not considered as a critical parameter for its function and transport until recently when it was reported to play a major role in determining the decay of cellular mRNAs and circRNAs This process has been termed as structure-mediated RNA decay (Fischer et al., 2020). Specifically, highly structured linear and circular RNAs are more prone to decay in the cell compared to those with lower structuredness (Fischer et al., 2020). Here, we report that structured RNA decay does have implications on the transport of RNAs into EVs wherein highly structured RNAs tend to be retained in the cell and not get packaged into EVs while less structured RNAs tend to get enriched into EVs. We applied our observations to publicly available datasets for circRNAs as well as linear RNAs and found that structuredness was consistently inversely correlated with EVs enrichment of RNAs. Furthermore, our data suggest that circularization of RNA leads to a decrease in its structuredness, and that this could explain the preferential enrichment of circRNAs in EVs over their cognate linear isoforms. Specifically, we found that most circRNAs have lower size-normalized structuredness than their linear isoforms and that the enrichment of circRNAs is independent of their linear isoforms. This selective enrichment may be due to the enhanced stability of circRNAs as they are less likely to be targeted by structure-mediated RNA decay pathway. Furthermore, it is likely that different RNA binding proteins recognize RNAs based on their structure instead of just the sequence motifs. In other words, the actual “zipcode” might lie in the secondary structure of RNA instead of its linear sequence.

Thus, we found that a combination of multiple factors contribute to the enrichment of circRNAs in EVs. The higher abundance of circRNAs in EVs could enhance their role in modulating gene expression in recipient cells, suggesting a preferential mechanism for EV-mediated transfer of regulatory RNAs as well as miRNAs which had been extensively reported in literature (Bao et al., 2018; Abels et al., 2019; Harmati et al., 2019). Our data indicate a that there is a selection process in the cell where RNAs of different classes either get sorted to EVs or are retained in the cell based on their cis elements (Figure 7). It is likely that this process is mediated by RBPs that recognize these elements along the RNA sequence and effect their transport (O’Grady et al., 2022). For instance, RBPs could bind to stem-loop or other structured parts of an RNA leading to its retention in the cell whereas those lacking such structured elements are free to bind other RBPs that facilitate their enrichment in EVs.

**Figure 7.**
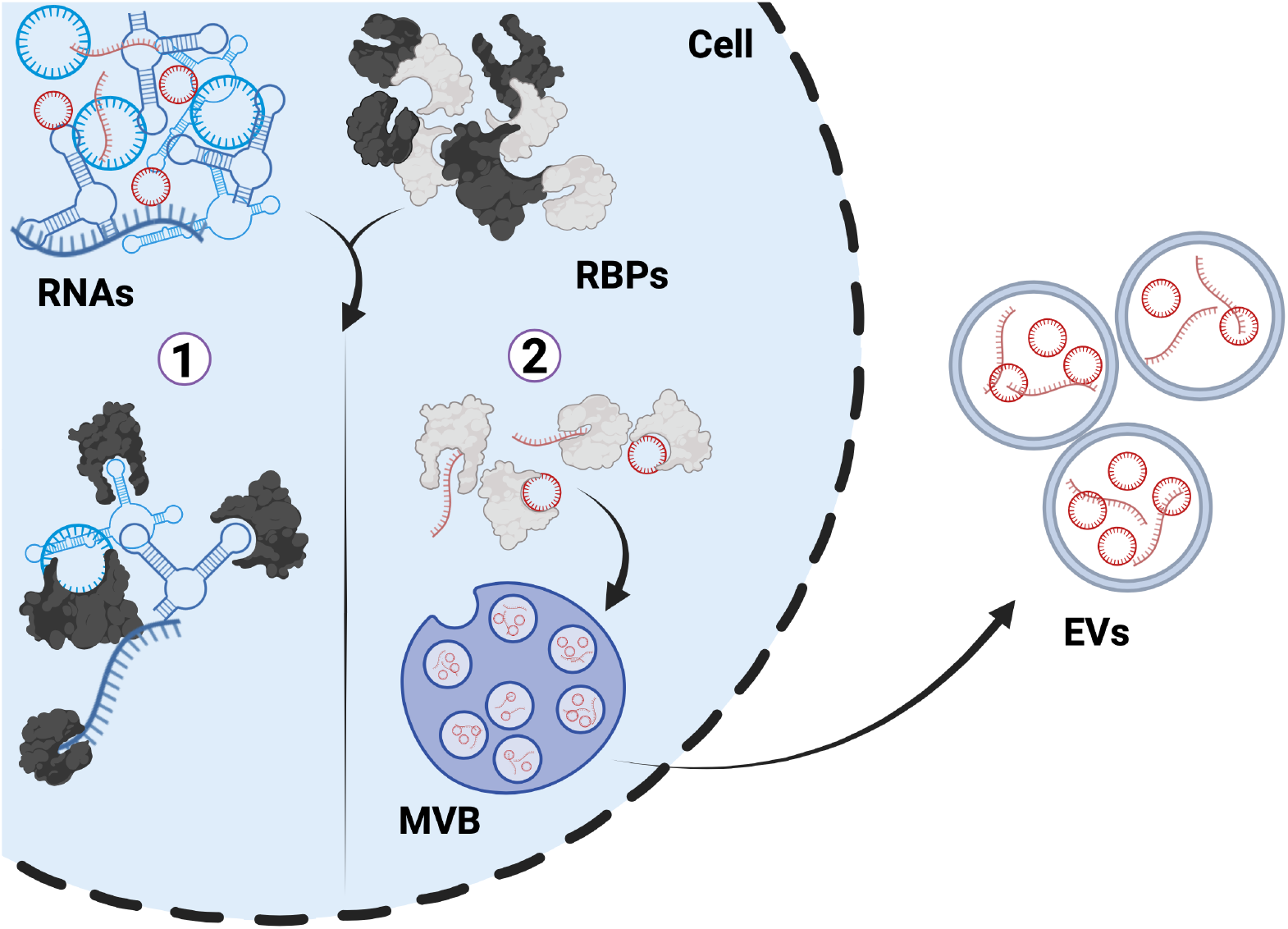
Model of RNA sorting into EVs: MVB = multivesicular body. Different colored proteins are depicting unique proteins that interact with high or low structured RNAs. Created with BioRender.com.

Cellular origin could impact RNA profiles in EVs and hence conducting such analysis using a few cell lines under specific conditions may not fully capture the diversity of RNA enrichment patterns in EVs (Vaka et al., 2023). For this reason, we sought to validate our results using existing publicly available databases such as exoRBase and circBase. In the case of circRNAs and mRNAs, we were able to validate most of the results from our datasets. In the case of lncRNAs, however, we observed less consistency with the results from our dataset. Only coding probability showed consistent results for lncRNAs in exoRBase. However, results from our dataset as well as in O’Grady et al. indicate that EVE lncRNAs have more exons than CR lncRNAs whereas there was no significant difference detected in exoRBase dataset (O’Grady et al., 2022). It is possible that different factors could be at play in different cell types or in response to various environmental conditions. Furthermore, although we have focused in this study on the cis elements of RNAs that are correlated with their enrichment in EVs, these effects are probably mediated through a network of RBPs in the cell. Ongoing work aims to identify RBPs that are potentially involved in RNA sorting into EVs and investigate the specific molecular mechanisms underlying these interactions. Functional validation experiments, such as RNA immunoprecipitation or knockdown studies, are then needed to confirm the role of these proteins in RNA packaging and to determine if other unidentified factors contribute to this process.

Differentiating reads originating from circRNAs and mRNAs in RNA-Seq data has been quite challenging. To investigate circRNAs, we utilized RNase R treatment for the samples in this study which is a 3’-5’ exoribonuclease that degrades linear RNAs with high processivity and, thereby, enriching circRNAs. However, RNase R’s activity is not 100% effective in degrading linear RNAs. Specifically, RNase R needs at least 7 unstructured nucleotides at the 3’ end of RNAs to be able to bind those RNAs (Vincent and Deutscher, 2006). Moreover, it was reported RNase R is interrupted by internal G-quadruplex structures (Xiao and Wilusz, 2019). Therefore, it remains difficult to accurately distinguish reads from linear and circular transcripts. Hence, we relied on back-splice junctions to count circRNAs for most of our analysis. Imaging approaches such as circFISH can be used to experimentally quantify the difference between circular and linear transcripts of a given gene (Koppula et al., 2022). While this definitely gives an underrepresented count of circRNAs in a sample, it lends more confidence to the conclusion of our analysis. This could explain the discrepancy between studies reporting circRNAs enrichment in EVs and those unable to detect such enrichment. The latter studies have likely relied solely on back splice junctions to quantify circRNAs enrichment (Pérez-Boza et al., 2018). Full-length RNA-Seq, and perhaps in conjunction with short-read sequencing, could be able to address this challenge and to further identify alternatively spliced circRNAs isoforms that share the same back splice junction (Xin et al., 2021; Kontos et al., 2023).

While our study sheds light on new factors influencing RNAs enrichment in EVs, several limitations need to be acknowledged. The methods used for EV isolation could introduce biases in the types of RNAs detected and EVs of different size could have different cargo (Tang et al., 2017; Barreiro et al., 2020). In this study, most of our EVs population were within the size range of small EVs (sEVs) (<200 nm) (Welsh et al., 2024). It would be interesting to see if these findings would be applicable to EVs of larger size ranges. Moreover, in some EVs isolation techniques, other extracellular components, such as lipoproteins or protein complexes could be co-isolated with EVs, which could confound the analysis of RNA content. For this reason, we included a size exclusion step in our EV isolation protocol to remove such confounding factors. Additionally, many of the steps in a RNA-Seq protocol can introduce biases, particularly for samples with low quantity or low quality of RNA which can eventually affect the comprehensiveness of RNA profile analysis (Shi et al., 2021). Further studies using a broader range of cell types and experimental conditions would be important to generalize these findings. Furthermore, our study primarily focused on the steady-state levels of RNA in EVs without considering the dynamics of RNA packaging and release. RNA content in EVs may change over time or in response to cellular or environmental stimuli, and these temporal dynamics were not captured in our experimental design. Longitudinal studies with time-course analysis would provide a more comprehensive understanding of how RNA cargo is selectively packaged into EVs under these different conditions. Addressing these limitations in future research will be critical to fully understanding the mechanisms governing RNA enrichment in EVs and their potential applications in therapeutics.

Our study provides insights into the factors that govern RNAs enrichment in EVs, emphasizing the selective nature of RNA packaging influenced by multiple cis elements. Ongoing work in our lab is focused on determining if RNA structure lays a role in intracellular RNA transport as well and to characterize RNA binding proteins that are involved in distinguishing EV cargo based on RNA structure and the underlying mechanisms for this selection process. Understanding these mechanisms is crucial for leveraging EVs in therapeutic applications, such as RNA delivery vehicles. Future work should focus on elucidating the detailed molecular mechanisms driving selective RNA packaging and enrichment and how these processes are altered in different pathological states. This could pave the way for more precise use of EVs in therapeutics applications.

## 4. Materials and Methods

### 4.1. Cell culture

DLD-1 cells (a generous gift from Leung lab (Johns Hopkins, MD, USA)), HEK-293T, and HeLa cells were cultured in DMEM (Millipore Sigma, St. Louis, MO, USA, D6429) supplemented with 10 % FBS (Millipore Sigma, St. Louis, MO, USA, F2442) and 1% penicillin/streptomycin (Millipore Sigma, St. Louis, MO, USA, P4333). All cells were grown in a humidified incubator at 37 °C and with 5% CO2. Lentivirus for creating stable cell lines expressing linear (Addgene 12154) and circular GFP (circGFP) (a gift from Guarnerio Lab) in 293T cells and then added to HeLa cells (Guarnerio et al., 2019). Successfully transduced cells were sorted using Fluorescence Activated Cell Sorting (FACS) (BD FACSAria III, San Jose, CA, USA).

### 4.2. EVs isolation and characterization (nanoFCM)

400 mL of Cell culture-conditioned medium (CCM) was spun down using ThermoFisher Sorwall WX80 centrifuge at 10,000 × g for 30 min at 4°C with max. acceleration and deceleration using Open-Top Thinwall Ultra-Clear Tubes (Beckman Coulter, 344058) and AH-629 swinging bucket rotor (K factor: 242). The subsequent supernatant from the spin was centrifuged again at 100,000 × g for 70 min at 4°C. Resulting pellet was resuspended in 1mL PBS and loaded onto a qEV1 Legacy 70 nm column (Izon) prewashed with Dulbecco’s phosphate buffered saline (DPBS, Gibco) and four 1-mL fractions were collected using automated fraction collector (Izon) with default settings. Fractions were combined and further concentrated with Amicon 15 Ultra RC 10 kDa filters (Millipore Sigma at room temperature (RT, approx. 22C) till final volume.

Flow NanoAnalyzer (NanoFCM, Inc) was used to measure the concentration and size of particles following the manufacturer’s instructions. Briefly, the instrument was calibrated separately for concentration and size using 250 nm PE- and FITC-fluorophore-conjugated Silica Nanospheres (NanoFCM, QS2502) and a Silica Nanosphere Cocktail #1 (NanoFCM, S16M-Exo), respectively. Samples were diluted in DPBS, and events were recorded for 1 minute. Using the calibration curve, the flow rate and side scattering intensity were converted into corresponding particle concentrations and size.

### 4.3. Western Blot

The volume of the final product normalized for all samples. 10 uL of the sample were mixed with 8 uL of 1x PBS and were lysed in 1x radioimmunoprecipitation assay buffer (RIPA, Cell Signaling Technology, 9806) for 30 minutes at RT. Lysates were heated at 100C for 5 minutes with Laemmli sample buffer (Bio-Rad, 1610747).

Samples then were subjected to SDS-PAGE using a 4% to 15% Criterion TGX Stain-Free Precast gel (Bio-Rad, 5678084), then transferred onto a PVDF membrane (Invitrogen, IB24001) using iBlot 2 semi-dry transfer system (Invitrogen) for 1 minute at 20 V, 4 minutes at 23 V, and 2 minutes at 25 V. Blots were incubated for one hour in PBST (PBS with 0.05% Tween-20 (BioXtra, P7949) and 5% Blotting Grade Blocker (Bio-Rad, 1706404) (PBST-Milk) and incubated overnight (approx.16 hours) in primary antibodies (CD63 BD Biosciences, 556019 1:1000 dilution, CD9 Biolegend, 312102 1:1000 dilution; Syntenin-1 Abcam, ab133267 1:1000 dilution; and GM-130 Santa Cruz, sc-55590 1:500 dilution). Membranes were washed three times in PBST-milk before incubation with species-specific HRP-conjugated secondary antibodies (m-IgGκ BP-HRP Santa Cruz, 616102 1:10000 dilution; or Mouse anti-Rb IgG HRP, Santa Cruz 2327 1:10000 dilution). SuperSignal West Pico PLUS Chemiluminescent Substrate (Thermo Scientific, 34580) was used for detection, and blots were visualized with an iBright 1500FL Imager (Thermo Fisher).

### 4.4. Transmission Electron Microscopy

Extracellular vesicle preparations (10 µl) were imaged as described in Huang et al. (2020) with a Philips CM120 instrument and an 8-megapixel AMT XR80 charge-coupled device (Huang et al., 2020).

### 4.5. RNA extraction, library preparation and sequencing

RNA extraction and total transcriptome sequencing were conducted as described in Huang et al (2024) (Huang et al., 2024). In brief, total RNA was extracted from cells with Trizol (Thermo Fisher, 15596018) or 100 µL EVs with Trizol LS (Thermo Fisher; 10296028). After homogenization, RNA was isolated using miRNeasy solutions (Qiagen, 217004) with Zymo-Spin columns (Zymo Research, C1003-50). RNA purity and concentrations were measured using Nanodrop (Thermo Scientific, Waltham, MA, USA, ND-2000).

RNAs from cells (2,000 ng) and EVs (960 ng) was incubated with 1 U/µg RNase R (Lucigen, RNR07250) at 37°C, 30 minutes and purified by RNA Clean&Concentrator-5 (Zymo Research, R1014). cDNA libraries (100 ng RNA, with/without RNase R) were made with “IDT for Illumina RNA UD” indices by Stranded Total RNA Prep Ligation w/Ribo-Zero Plus (Illumina, 20072063). Yield/size distribution were assessed by Fragment Bioanalyzer (Agilent, DNA 1000, 5067-1505). Multiplexed libraries were equally pooled to 2.5 nM and sequenced by NovaSeq 6000/S1 Reagent Kit version 1.5 (Illumina, 300 cycles; 20028401).

### 4.6. RT_-q_PCR and Sanger sequencing

Total RNA was extracted from cells or EVs by lysis in Trizol (Sigma, St. Louis, MO, USA, T9424) using the phenol-chloroform method, following the manufacturer’s protocol. RNA purity and concentrations were measured using Nanodrop (Thermo Scientific, Waltham, MA, USA, ND-2000). Equal concentrations of RNA from cells and EVs were used for cDNA synthesis using iScript Reverse Transcription Supermix (Bio-RAD, Hercules, CA, USA, 1708841), and gene expression was analyzed using iTaq Universal SYBR Green Supermix (Bio-RAD, Hercules, CA, USA, 1725124) according to the manufacturer’s protocol. All of the experiments were performed in triplicates. CT values were used to calculate fold change (2-ΔΔCT) using Actin (ACTB) RNA as control (Schmittgen and Livak, 2008). The p-values were obtained using Student’s t-test. PCR products were visualized by agarose gel electrophoresis to confirm amplification and assess fragment size. Circular GFP (circGFP) amplicons from cellular and EVs fractions were purified using QIAEX II Gel Extraction Kit (Qiagen, Hilden, Germany, 20021) and sent for Sanger sequencing (Azenta Life Sciences, South Plainfield, NJ, USA). All of the primers used are provided in Supplementary Table S1.

### 4.7. CircRNAs analysis

For RNA-Seq data: Quality control of RNA-Seq data was performed using FastQC and MultiQC (Ewels et al., 2016). Data were aligned to human genome (hg38) using STAR 2.7.10b and circRNAs were identified using CircExplorer2 2.3.8 (Zhang et al., 2016; Dobin et al., 2013). CircRNA exon sequence coordinates were obtained based on all exon sequences in UCSC known gene annotations (v40) that overlap the circRNAs coordinates reported by circExplorer2 using GenomicFeatures 1.54.4 R package (Lawrence et al., 2013). To ensure the accuracy of circRNAs annotations, both start and end positions of circRNAs had to match an exon boundary in the annotations. Next, circRNAs sequences were extracted from human genome (hg38) using BSgenome 1.70.2 R package. When comparing the enrichment of circular and linear RNAs in EVs, Kallisto 0.44.0 was used to align the reads from RNase R-treated samples to human genome (hg38) and these counts were compared to mRNA counts obtained from mock-treated samples (Bray et al., 2016). For circRNAs in ExoRBase 2.0 and CircBase, transcripts with no stop codons were first discarded from UCSC known gene annotations (v40) then exons that intersect with circRNAs coordinates were obtained using GenomicFeatures 1.54.4 R package) (Glažar et al., 2014; Lai et al., 2022; Lawrence et al., 2013). Next, sequences were extracted similarly. For circRNAs quantification, number of forward-splice (fsj) and back-splice (bsj) junction reads were obtained using CIRIquant 1.1 with default parameters (Zhang et al., 2020). Then, the ratio of circular/linear RNAs were calculated as follows: 2 ^*^ bsj / (2 ^*^ bsj + fsj).

### 4.8 Linear RNA analysis

For all linear RNA analysis, control (no RNase R treatment) samples were used. First, Gencode v32 annotations was obtained and divided into three categories: mRNAs containing protein-coding RNAs, lncRNAs containing long non-coding RNAs, and others containing all other transcripts. Next, Gencode v32 annotations were then used to further separate mRNA transcripts into 5’ UTR, CDS, and 3’UTR using GenomicFeatures 1.54.4 R package, and their sequences were obtained from human genome (hg38) using BSgenome 1.70.2 R package (Lawrence et al., 2013). Individual indexes were built for the three linear RNA groups from the extracted sequences using Kallisto 0.44.0 and reads were then aligned to those sequences (Bray et al., 2016). Counts were imported using tximport 1.30.0 R package and analyzed for differential expression using DESeq2 1.42.1 R package (Love et al., 2014). ExoRBase sequences were obtained by first matching the gene ID from the database to that in Gencode v32 annotations and then extracting the sequences from human genome (hg38) using BSgenome 1.70.2 R package.

### 4.9. Differential expression analysis

Differential expression was performed using DESeq2 1.42.1 R package with default parameters (Love et al., 2014). RNAs were labeled as EVs-enriched (EVE) if their adjusted p-value (FDR) < 0.05 and fold change > 0, or as cell-retained (CR) if FDR < 0.05 and fold change < 0, or as not significantly different (NS) otherwise.

### 4.10. Secondary structure analysis

RNAfold 2.4.13 from ViennaRNA package was used to calculate the minimum free energy (-ΔG) of RNAs which was then inversed and normalized to the length of RNAs (-ΔG/nt) (Lorenz et al., 2011). The “--circ” option was used for circRNAs or to calculate (-ΔG/nt) of linear sequence in circularization experiment. Due to size limitation of RNAfold, sequences bigger than 32,767 nt were not included in the analysis.

### 4.11. Protein abundance

Cells were collected and rinsed three times with cold PBS, and then lysed with SDS buffer (5% SDS, 100mM Tris-HCl, pH=8). The lysate was digested using E3filter (CDS Analytical, Oxford, PA) as described previously (Martin et al., 2024). The digests were desalted using C18 based StageTips, dried in SpeedVac, and stored under -80 °C before further analysis. The peptides were analyzed on an Ultimate 3000 RSLCnano system in conjunction with an Orbitrap Eclipse mass spectrometer and FAIMS Pro Interface (Thermo Scientific) following a method published previously (Martin et al., 2024). For data-independent acquisition (DIA) and FAIMS settings, we adapted a protocol reported in a previous paper (Elsayyid et al., 2024). The DIA mass spec data were processed using Spectronaut software (version 19.1) with most of the default settings (Bruderer et al., 2015). Briefly, a library-free DIA analysis workflow with directDIA+ and an UniProt human protein database (82,861 sequences; version July 2024) were used. The settings for Pulsar and library generation include: Trypsin/P as specific enzyme; peptide length from 7 to 52 amino acids; allowing 2 missed cleavages; toggle N-terminal M is true; Carbamidomethyl on C as fixed modification; Oxidation on M and Acetyl at protein N-terminus as variable modifications; FDRs at PSM, peptide and protein level all set to 0.01. Next, the output intensity-based absolute quantification (iBAQ) values from the protein quantity report were correlated with TPM values for mRNAs from Kallisto 0.44.0 (Bray et al., 2016).

## Supplementary Materials

Supplementary figures, Supplementary table 1.

## Data Availability Statement

Sequencing data were deposited in GEO (GSE279376).

## Acknowledgments

We would like to thank the members of Batish and Parashar Laboratories for their discussion and support.

## Ethics approval statement

N/A

## Patient consent statement

N/A

## Permission to reproduce material from other sources

N/A

## Clinical trial registration

N/A

## Author Contributions

AA: Investigation, data curation, analysis, methodology, article writing, review, editing. YH: Resources, methodologies and data curation, review, editing. OG: Resources, methodologies and data curation, review, editing. YY: Resources, methodologies and data curation, review, editing. KWW: supervision, review, editing. VP: supervision, funding acquisition, review, editing. MB: conceptualization, supervision, validation, funding acquisition, project administration, article writing, review, and editing.

## Supporting Information

**Extended Data Figure 1.**
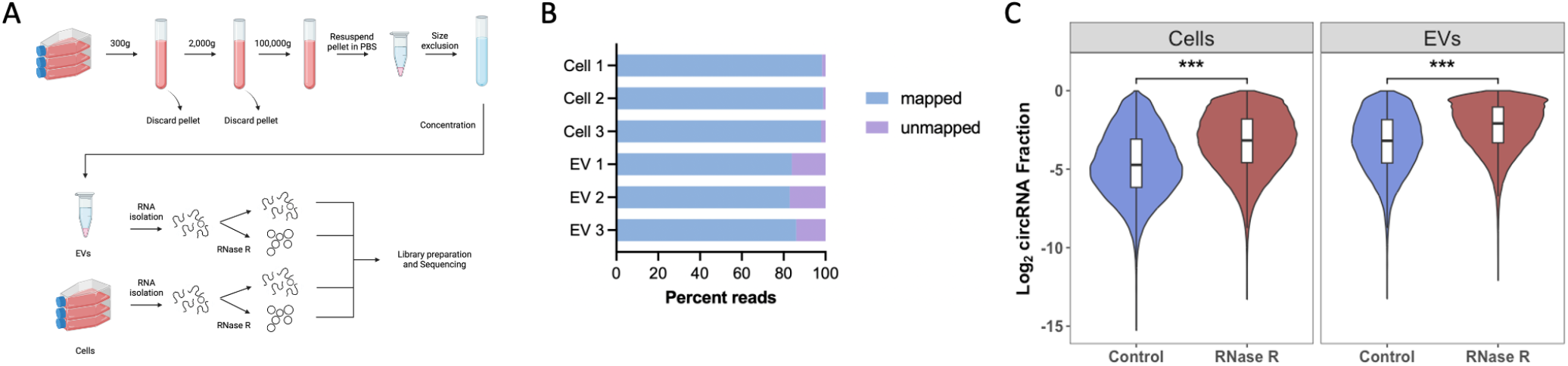
RNA-Seq workflow and data quality: **(A)** Workflow of EVs isolation and RNA-Seq. **(B)** Percentage reads mapped to human genome (hg38) or unmapped from RNA-Seq for three EV and three cell samples. **(C)** Log2 circular/linear RNA ratio in control and RNase R-treated samples in cells and EVs.

**Extended Data Figure 2.**
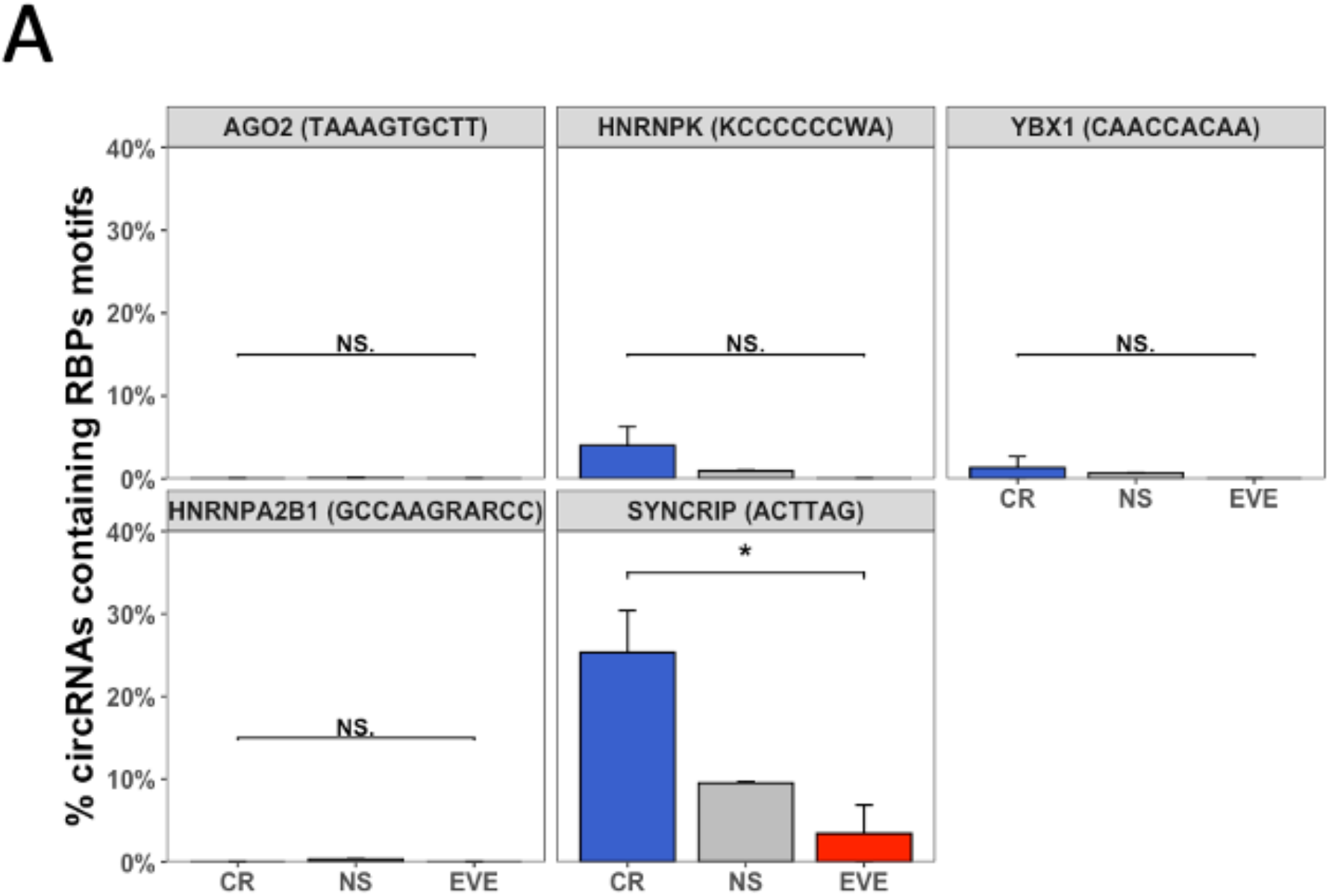
Role of cis elements in EVs enrichment of circRNAs: **(A)** Percentage of EVs enriched (EVE), cell retained (CR), or circRNAs with no significant difference (NS) that contain RBPs motifs. Motifs of RBPs are shown in parenthesis. EVE = EVs enriched, NS = not significant. CR = cell retained. P-values were calculated using Mann-Whitney’s test, *p < 0.05, **p < 0.01, ***p < 0.001.

**Extended Data Figure 3.**
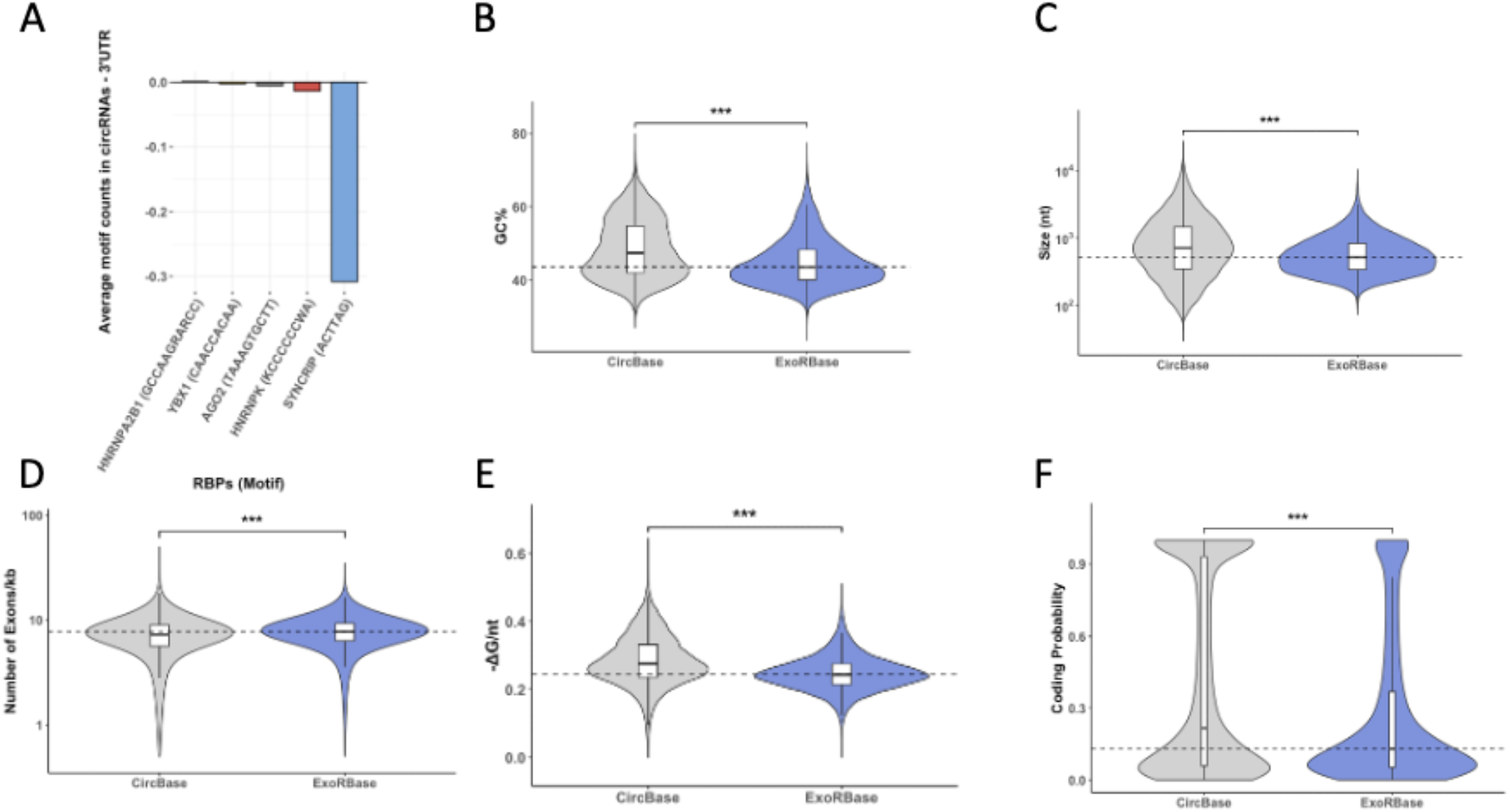
Validation of circRNAs cis elements from databases: **(A)** Average motif count difference between circRNAs in circBase and the 3’ UTR of their linear counterparts for five RBPs. Motifs of RBPs are shown in brackets. **(B-F)** Violin plot of GC%, size, number of exons per kilobase of transcript, structuredness, and coding probability, respectively, of circRNAs in circBase or exoRBase. P-values were calculated using Mann-Whitney’s test, *p < 0.05, **p < 0.01, ***p < 0.001.

**Extended Data Figure 4.**
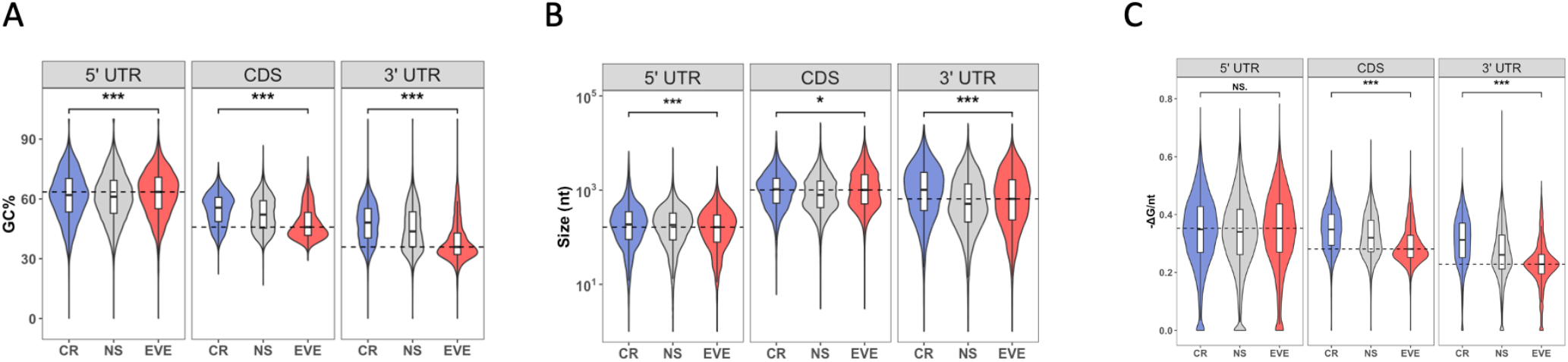
Evaluation of cis Elements Across 5’UTR, CDS, and 3’UTR Regions of mRNAs: **(A)** Violin plot of GC% in 5’ UTR, CDS and 3’ UTR of mRNAs separated based on their enrichment into EVE, NS and CR RNAs. F. Similar to R but for size. **(B-C)** Similar to A but for size and structuredness, respectively. EVE = EVs enriched, NS = not significant. CR = cell retained. P-values were calculated using Mann-Whitney’s test, *p < 0.05, **p < 0.01, ***p < 0.001.

**Extended Data Figure 5.**
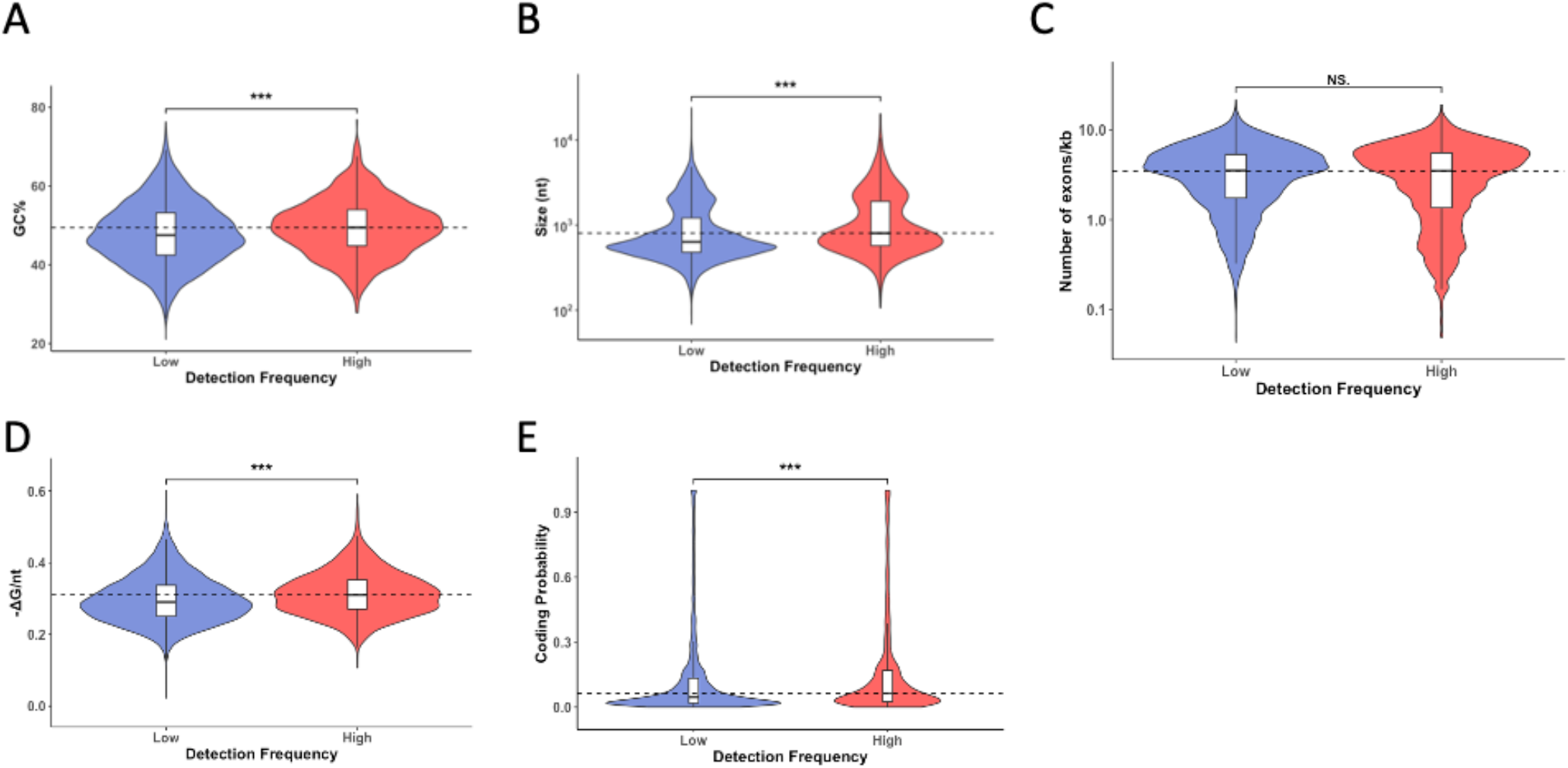
Validation of lncRNAs cis elements from databases: **(A-E)** Violin plot of GC%, size, number of exons per kilobase of transcript, structuredness, and coding probability, respectively, of exoRBase lncRNAs with low or high detection frequency. P-values were calculated using Mann-Whitney’s test, *p < 0.05, **p < 0.01, ***p < 0.001.

**Extended Data Figure 6.**
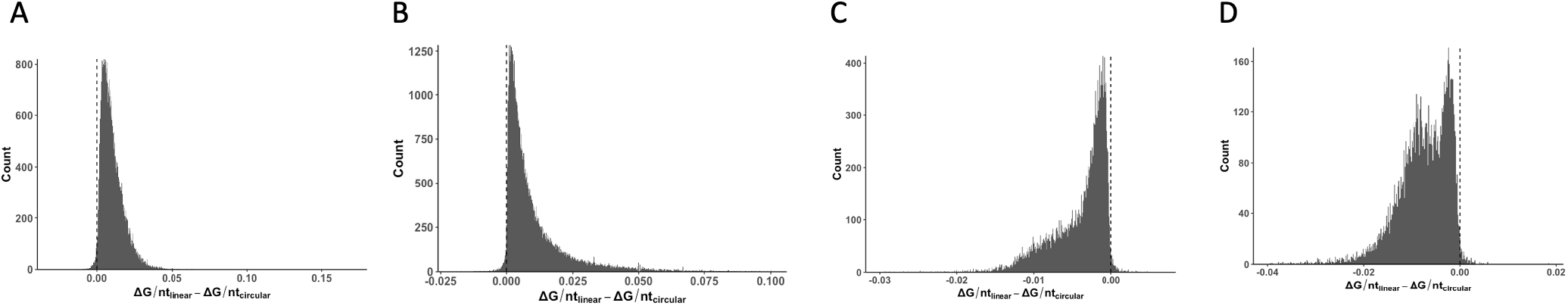
Validation of the role of structuredness in databases: **(A-B)** Change in structuredness of after linearization of circRNAs sequences in exoRBase and circBase, respectively. **(C)** Change in structuredness of after circularization of mRNAs sequences in exoRBase. **(D)** Similar to C but for lncRNAs.

## Supplementary Table

List of primers used in the study

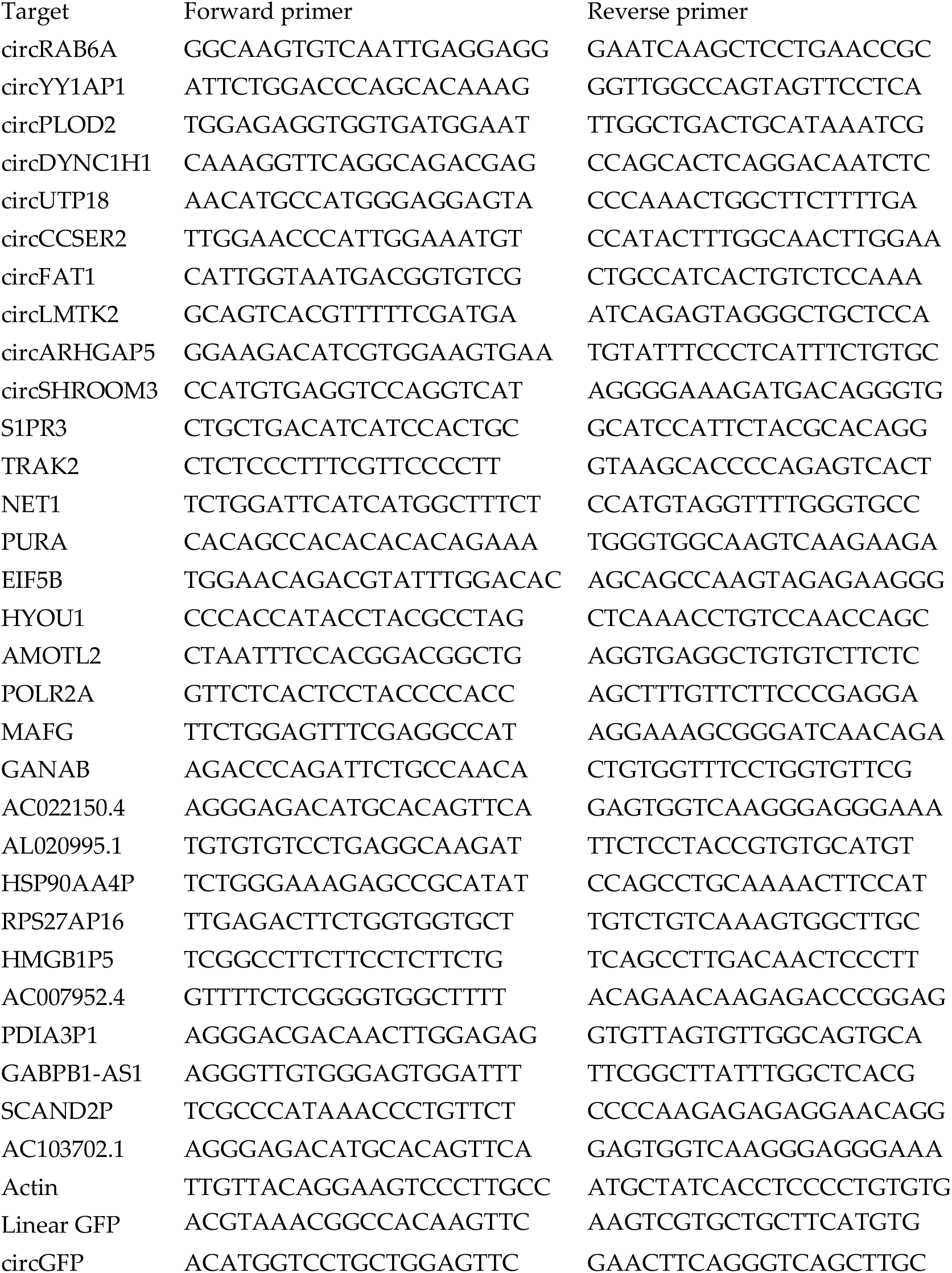

